# Rummagene: Mining Gene Sets from Supporting Materials of PMC Publications

**DOI:** 10.1101/2023.10.03.560783

**Authors:** Daniel J. B. Clarke, Giacomo B. Marino, Eden Z. Deng, Zhuorui Xie, John Erol Evangelista, Avi Ma’ayan

## Abstract

Every week thousands of biomedical research papers are published with a portion of them containing supporting tables with data about genes, transcripts, variants, and proteins. For example, supporting tables may contain differentially expressed genes and proteins from transcriptomics and proteomics assays, targets of transcription factors from ChIP-seq experiments, hits from genome-wide CRISPR screens, or genes identified to harbor mutations from GWAS studies. Because these gene sets are commonly buried in the supplemental tables of research publications, they are not widely available for search and reuse. Rummagene, available from https://rummagene.com, is a web server application that provides access to hundreds of thousands human and mouse gene sets extracted from supporting materials of publications listed on PubMed Central (PMC). To create Rummagene, we first developed a softbot that extracts human and mouse gene sets from supporting tables of PMC publications. So far, the softbot has scanned 5,448,589 PMC articles to find 121,237 articles that contain 642,389 gene sets. These gene sets are served for enrichment analysis, free text, and table title search. Users of Rummagene can submit their own gene sets to find matching gene sets ranked by their overlap with the input gene set. In addition to providing the extracted gene sets for search, we investigated the massive corpus of these gene sets for statistical patterns. We show that the number of gene sets reported in publications is rapidly increasing, containing both short sets that are highly enriched in highly studied genes, and long sets from omics profiling. We also demonstrate that the gene sets in Rummagene can be used for transcription factor and kinase enrichment analyses, and for gene function predictions. By combining gene set similarity with abstract similarity, Rummagene can be used to find surprising relationships between unexpected biological processes, concepts, and named entities. Finally, by overlaying the Rummagene gene set space with the Enrichr gene set space we can discover areas of biological and biomedical knowledge unique to each resource.

## Introduction

The introduction of omics technologies has gradually moved biological and biomedical research from studying single genes and proteins towards studying gene sets, clusters of genes, molecular complexes, and gene expression modules^1^. Many biomedical and biological research studies produce and publish gene and protein sets. For example, differentially expressed genes and proteins from transcriptomics and proteomics assays, genes associated with genomic variants identified to be relevant to a phenotype, gene knockouts associated with a cellular or an organismal phenotype, target genes of transcription factors as determined by ChIP-seq experiments, proteins identified in differential phosphoproteomics, proteins identified in a complex from immunoprecipitation followed by mass-spectrometry studies, genes associated with a cellular phenotype from CRISPR screens, and many more types of sets can be generated. These gene sets are highly valuable but not often reused. This lack of reuse is partially because there are no standards for submitting gene sets in publications, and there are no community repositories for depositing gene and protein sets. As a result, the potentially useful information about gene sets is buried in supporting material tables stored as PDF, Excel, CSV, or Word file formats. Since general and domain specific search engines do not index the contents of such supporting materials, there is no way to search through these tables. These supporting tables are not indexed by search engines because most search engines can only deal with free text and are not capable of parsing data tables.

Named entity recognition methods have been widely applied to biomedical and biological publication text, but not yet to extract gene sets from supporting tables. Manual gene set annotations and extraction of gene sets from publications has been achieved, but it is time consuming, labor intensive, and requires domain expertise. Most such efforts miss many relevant studies. For example, to create the ChIP-x Enrichment Analysis (ChEA) resource we manually extracted gene sets from supporting materials of ChIP-seq studies^2,3^. While the ChEA database achieved great success, it is difficult to maintain. Efforts such as ReMap^4^, Recount^5^, and ARCHS4^6^ aim to address this challenge by uniformly reprocessing all the raw data available from community repositories to recompute gene sets from published studies, but such efforts rely on the existence of community repositories and uniform data collection standards. Another effort to automate the extraction of gene sets from publications is Pathway Figure Optical Character Recognition (PFOCR)^7^. PFOCR automatically extracts pathways from publications by scanning pathway diagrams. However, surprisingly, as far as we know there are no publications, databases, or community repositories that contain extracted gene sets from supporting materials of scientific biomedical research publications. Rummagene is a web-based software application that serves hundreds of thousands of gene sets extracted from publications listed on PubMed Central (PMC). It contains a softbot that scans supporting materials of publications listed on PMC to keep the resource consistently updated. The Rummagene website provides the ability to search the corpus of gene sets by an input gene set query, a PMC free text search, and a table title search. To understand the statistical patterns within the Rummagene corpus, we performed various exploratory analyses, as well as demonstrate how this rich resource of organized biological knowledge can be used for specific applications.

## Methods

### Crawler to extract gene sets from publications listed in PMC

The PMC Open Access Subset^8^ contains millions of journal articles available under license terms that permit reuse. Additionally, PMC provides uniformly structured bundles that can be retrieved in bulk over FTP. An index file contains a tabular listing of all PMCIDs represented with a pointer to the compressed bundle corresponding to that PMCID. Each bundle has a PDF of the paper, an XML document containing structured metadata about the paper, figures, and supplemental material files. First, the index file is downloaded, a job is then submitted for each paper. The job downloads and extracts the archive, and every file in the bundle is processed by one of several table-extractor-functions, selected based on the file extension. These extractor functions include support for Excel, CSV, TSV, and inferred separator loading of TXT files, as well as a PDF table extractor based on Tabula-Py, and an extractor function that grabs tables from the structured metadata provided by PMC about the tables in the articles. For each supporting materials table that is extracted, every column in the table is considered. The extractor function attempts to map all unique strings to gene symbols. Mapping may be direct, through some synonym, or identifier. Any column where more than half of the strings can be successfully mapped to a valid gene symbol using NCBI’s Gene Info^9^ file for *Homo sapiens* are retained. In other words, all columns passing this filter become a gene set in the Rummagene gene set library. The term describing the gene set is made of the PMCID, the file name in the bundle, the Excel spreadsheet name or the XML table label, the column’s first cell, and additional sequential numbers that are added to the term to make it unique if needed. The original items in each table column that pass the filter are preserved, but genes are included only if they can be mapped to official symbols. In addition to filtering out columns with too few mapped genes (<5), columns with too many mapped genes (>2000) are ignored. This is because these are likely to contain gene sets that cover all measured genes and not a subset of identified genes with a potentially unique function. This pipeline produces a large gene matrix transpose (GMT) file which can be added to incrementally. The pipeline is designed to continue where it left off when it is re-run. It is set to run weekly to extend the database with any new publications that are added to the PMC Open Access database. The new entries to the GMT are stored in the Rummagene database to be accessed from the web-based application.

### Search engine implementation

The large size of the Rummagene gene set library requires special implementation of an algorithm that can quickly compare the input gene set to all the gene sets in the Rummagene database. Besides a fast algorithm that can compare the input set to all other sets, efficient storage of the gene sets is needed as well as sufficient hardware. To enable a fast gene set search, a Rust-powered REST API was implemented. The algorithm first initializes several in-memory data structures: 1) a background sorted set of genes; 2) the index of each gene saved in a hashmap mapping where each gene is mapped to a 32-bit unsigned integer (U32) index; 3) the gene set IDs stored as UUIDs; and 4) a hashset of mapped genes using the Fowler–Noll–Vo (FNV) hash function. These data structures are created by querying the database with Rust. When the user presses the search button, the queried gene sets are forwarded to the API. After ensuring that the index is initialized, the code maps the user submitted gene set to a U32 hash set. It then computes the intersections between the user’s gene set and the gene sets in memory and performs the Fisher’s exact test using the identified overlap. Parallel processing with Rayon Rust^10^ is employed to further speed up this process. Once completed, Benjamini-Hochberg adjusted p-values are computed. Next, the results are sorted by p-value, temporarily cached, and returned. The gene sets in Rummagene are stored in a Postgres database^11^. A function in the Postgres database is responsible for mapping the gene symbols to UUIDs before passing them to the Rust API to obtain results. These returned results can be joined by ID with the gene sets and genes in the database to facilitate further filtering. In this way, the use of an API is transparent to the front-end which queries the database with PostGraphile powered GraphQL. When a batch of new gene sets are added to the database, a new background is constructed with the complete set of genes in the database. At that time, the API is called to prepare the new background prior to removing the old background.

### Extracting functional terms from column titles

To assess the contents of the extracted gene sets, the column titles for each table were examined to identify a variety of functional terms. Supplementary table titles often include DOI and other identification information, thus these were ignored when conducting this analysis. After separating column titles in each gene set, column titles were split on dashes, underscores, and periods. In identifying human genes, each resulting string was examined to assess if it was an NCBI Entrez^12^ approved gene symbol or a listed synonym. All gene synonyms were subsequently converted to their official symbol. Although genes can be represented with integer identifiers, strings only containing numbers were ignored after manual examination identified many of these as artifacts. Additionally, strings containing S succeeded by an integer were ignored considering the vast majority of these refer to the supplemental table number. Transcription factors and kinases were subsequently identified from the extracted gene symbols. To identify gene sets that may represent signatures, the strings ‘up’, ‘down’, and ‘dn’ were searched for in the split column titles. In order to identify tissues, cell types and cell lines present in the column titles, the Brenda Tissue Ontology (BTO)^13^ official terms and synonyms were extracted and exact matches were identified. Terms containing multiple BTO terms were hyphenated to capture, for instance, a cell type from a specific tissue.

### Visualization of the kinase and TF gene set libraries

For each extracted gene set, IDF vectors were computed using the Scikit-learn^14^ Python package using the set of all included genes as the corpus. Using the Scanpy^15^ Python package, Uniform Manifold Projection (UMAP)^16^ plots for different categories of gene sets were then generated from the IDF vectors and clusters were automatically computed using the Leiden algorithm^17^. To visualize broad patterns across the data, each point representing a gene set was colored based on the cluster, associated PMCID, and associated kinase or transcription factor, where applicable.

### Benchmarking transcription factor and kinase enrichment analyses

Consensus transcription factor and kinase gene set libraries were created by taking the union of all terms containing a given transcription factor or kinase in the column name. Benchmarking datasets were sourced from ChEA3^2^ for transcription factors and from KEA3^18^ for kinases. To benchmark enrichment analysis performed with the constructed consensus gene set libraries, the rank of each transcription factor/kinase was identified using the Fisher’s exact test p-value for each term in each benchmarking dataset. The cumulative distribution function of these ranks, D(r), was assessed. If the transcription factors or kinases do not tend to rank high or low, then it would be expected for D(r) to be equal to the uniform distribution r. Thus, D(r) – r can be used to represent significant deviations from zero and to assess the accuracy of the constructed gene set libraries. Kolmogorov-Smirnov tests for goodness of fit were used to evaluate the null hypothesis D(r) = r using the SciPy^19^ Python package.

### Topic modeling

To identify the predominant topics associated with gene sets in the Rummagene database, the abstracts of each paper contributing at least one gene set were assembled from the PMC bulk download. The text contained within the *<abstract>* tags was concatenated. Papers containing no abstracts were excluded from the analysis. Each abstract was then tokenized, stop words were removed, and lemmatized using the Python package Natural Language Toolkit (NLTK)^20^. The LdaModel class of Python package Gensim^21^ was then used to identify nine topics with a chunksize of 100 over 10 passes. The number of topics was chosen manually by observing the separation of topics given different sets of parameters. Word counts and word importance were extracted from the model for each of the nine topics. The abstracts were visualized in topic space using the vectors produced by the Latent Dirichlet Allocation (LDA) model^22^ for adherence of each paper to each topic using t-SNE^23^.

### Similar gene set pairs that are distant in abstract space

The preprocessing of publications’ abstracts followed the same procedure as in topic modeling where abstracts were first extracted from the PMC bulk download, then cleaned of stopwords and lemmatized using the NLTK^20^ Python package. Abstracts were then converted to word counts using the count vectorizer and subsequently fit to term frequency -inverse document frequency (TF-IDF) vectors using the Scikit-learn^14^ Python package. The cosine similarity of each paper abstract to all other abstracts was then assessed using the Scikit-learn pairwise linear kernel metric based on the computed TF-IDF vectors. Any pairs of papers with zero cosine similarity of their abstracts were retained, and for each gene set extracted from each of these papers, the Fisher’s exact test was performed to assess whether a significant overlap was present. Pairs with identical gene sets were excluded. The pairs were further filtered to only include overlaps of more than 50 genes. Additionally, to assess novelty of the recovered pairs, the percentage of their overlapping genes with ‘sticky proteins’ identified in analysis of protein-protein interactions^24^ were used. To assess the amount of highly cited genes, present in the overlapping genes of gene set pairs, the top 500 most cited genes according to GeneRIF^12^ were used. Additionally, to determine the amount of highly expressed genes present in the overlapping genes of gene set pairs, the top 500 most highly expressed protein coding genes were sourced based on mean expression across 5,000 random samples from ARCHS4^6^. In the analysis of gene set pairs including a gene or disease in the column title, only the top 10,000 most significant pairs with <10% ‘sticky proteins’ were included. To identify disease names in column titles of the gene set pairs, DisGeNet^25^ disease terms were used and gene names were identified using NCBI gene^12^ mappings. The OpenAI API chat completion module using the GPT-4 model was utilized to hypothesize the connection between the pairs of gene sets from this subset.

### Gene function predictions

50,000 gene sets were randomly selected from Rummagene and filtered for sets with less than 2000 genes. For all human genes, we formed a matrix *A* where *A*(*i, j*) = 1 if gene *i* is a member of term *j* and 0 otherwise. Then the co-occurrence matrix *Φ* = *A* · *A*^*T*^. As previously described^26^, the co-occurrence probability between two genes:

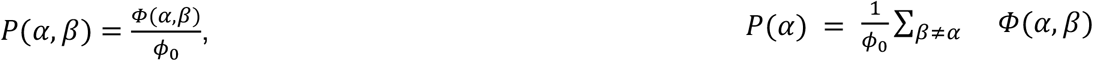

where *ϕ*_0_ is the total number of co-occurrences, and the marginal probability 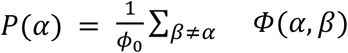.

The cosine similarity, Jaccard index, and normalized pointwise mutual information (NPWMI) for each pair of genes were then calculated as follows:

- *Cosine*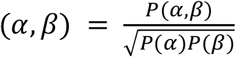
- *Jaccard*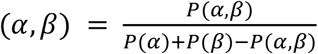
- *NPWMI*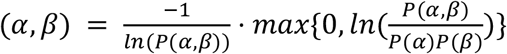

The NPWMI is a value between 0 and 1, where a larger value indicates the two genes co-occur with greater probability than expected by random chance^27^. Four libraries were used to benchmark gene function prediction: GO Biological Process (2023), GWAS Catalog (2023), MGI Mammalian Phenotypes (2021), and Human WikiPathways (2021). The distance of each gene to each term in the library was computed as the average distance of the gene to each constituent gene in the term gene-set. Suppose *L* is a matrix where *L*(*i, j*) = 1 if gene *i* is a member of term *j* in the library *L*, and 0 otherwise. Let *D* be the similarity matrix as described above, where the diagonal is set to 0. The gene-term association matrix 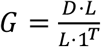 where the division is elementwise. Each entry *G*(*i, j*) is then the mean similarity of gene *i* to all the genes in gene set *j*. The matrix *G* can then be used to predict membership of gene *i* in any gene set. ROC curves and AUC values for each term in the library were computed using the Python sklearn.metrics module^14^.

### Comparing the Rummagene gene set space to the Enrichr gene set space

All the gene set libraries in Enrichr were assembled and processed together with the Rummagene gene sets so they can be projected into the same two-dimensional space. First, all genes were mapped to their official NCBI gene symbols for *Homo sapiens* or filtered out. Gene sets were then converted into vectors with values corresponding to the inverse document frequency (IDF)^28^. Truncated Singular Value Decomposition (Truncated SVD)^29^ was then used to reduce the dimensionality of the IDF vectors to the 50 largest singular values. A UMAP^23^ with the default settings was then used to embed all samples into two dimensions. Finally, to better position the visualization, we computed the mean and standard deviation of the embedding dimension axes and show the bulk of the samples that are within 1.68 standard deviations from the mean.

## Results

### Descriptive statistics

The initial version of Rummagene contains 642,389 gene sets extracted from 121,237 articles. These 121,237 articles are identified as containing gene sets from 5,448,589 scanned PMC articles. The distribution of the occurrence of genes in gene sets is not even. Some genes are found in many sets, but most genes are members of few sets (Fig. 1A). At the same time, most identified gene sets have less than one hundred genes in each set (Fig. 1B). While most publications only contributed to the Rummagene collection one or two gene sets, there are few publications that contributed a few hundred sets (Fig. 1C). Over the years, more and more gene sets are found in publications (Fig. 1D). In fact, in the past four years, publications included many more sets compared to sets identified in the 30 years between 1988 and 2018. Since 2005, the average length of gene sets jumped from less than 20 genes in each set to ∼150 genes in each set (Fig. 1E). This is likely due to the introduction of omics technologies and publications reporting gene sets identified from such studies. By projecting the gene set content into two dimensions with UMAP^16^, we see that, on average, short gene sets contain genes that are more commonly studied (Fig. 1F-G). While this is a general trend, some genes occur in many sets but are less commonly studied (Fig. 1H). This presents an opportunity to identify genes that are likely serving critical biological roles but are less explored.

**Fig. 1.**
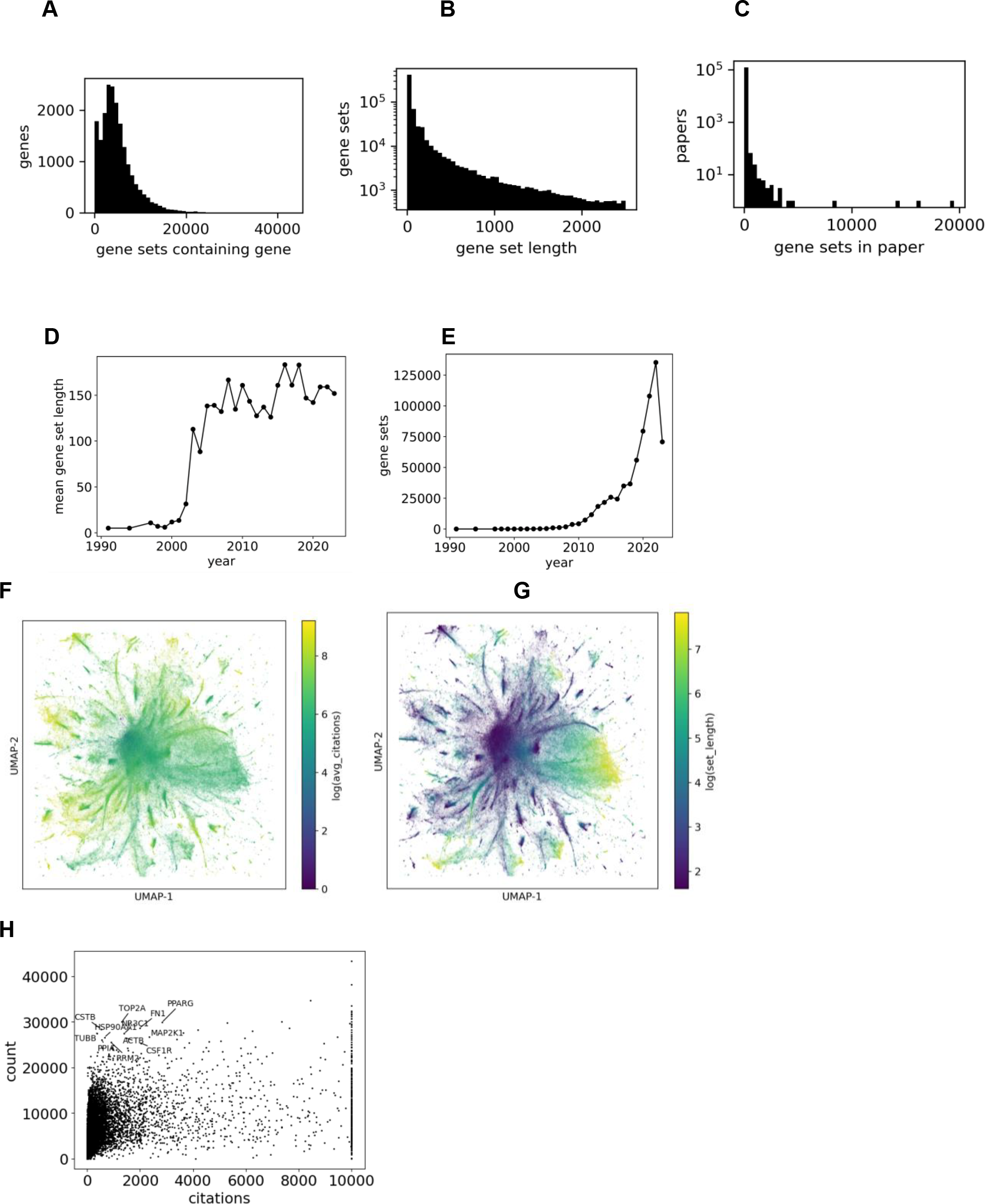
Distributions of the genes and gene sets in the Rummagene database. **A**. Distribution of genes in gene sets; **B**. Distribution of gene set lengths; **C**. Distribution of gene sets per paper; **D**. Gene set contribution per year. **E**. Average gene set length per year. **F**. Projection of the Rummage gene set into UMAP space where gene sets are colored by their average citations per gene. **G**. Projection of the Rummage gene set into UMAP space where gene sets are colored by their length. **H**. Scatter plot of all human genes with their overall citations vs. membership in Rummagene sets.

### Annotated collections of themed gene set libraries

For the collection of 642,389 gene sets we can identify subsets of gene sets associated with specific biological themes such as sets related to kinases, transcription factors, cell types, cell lines and tissues. Such themed gene sets can be used for specific enrichment analysis tasks such as kinase enrichment analysis^18^, transcription factor enrichment analysis^2^, and cell type identification for scRNA-seq. Producing such subsets of gene sets can be done by simply searching the table titles for terms that match named entities such as protein kinases or transcription factors. Indeed, we identified 4525 gene sets that contain named human kinase, and 8078 gene sets that contain named transcription factors in the table titles. 444 kinase names and 1121 transcription factor names are unique in these collections of gene sets (Fig. 2A). Similarly, we identified 4443 gene sets that contain named cell lines, and 6268 gene sets that contain cell types or tissues in table titles, with 450 and 670 unique terms, respectively (Fig. 2B, Table 1). In addition, 5560 sets had the term “down” and 6677 had the term “up” in their table titles (Fig. 2C). These sets likely contain up and down regulated genes from gene expression signatures. A large portion of the identified gene sets contain gene names in their titles. Specifically, 97,478 table titles contain human gene symbols or synonyms (Fig. 2C). For the subset of terms containing known transcription factors, UMAP plots were generated from the IDF vectors for all gene sets in the subset, and points representing different gene sets were colored by both the PMCID of the original publication, and by the associated transcription factor. We found that these gene sets tend to cluster by transcription factor (Fig. 2D) even when they are derived from different publications (Fig. 2E). We also applied the same process to generate UMAP plots for the subset of terms containing known kinases, and similarly saw that these gene sets clustered by kinase (Fig. 2F) across different PMCIDs (Fig. 2G). Next, we aimed to assess whether the kinase and transcription factor gene set libraries created from Rummagene contain useful information for performing gene set enrichment analysis. To achieve such an assessment, we queried each gene set from the Rummagene kinase and transcription factor libraries against corresponding kinase and transcription factor libraries created from multiple sources^2,18^. We observe a significant recovery of the correct kinases and transcription factors with all libraries with best agreement with PTMsigDB^30^ for kinases and ChEA 2022^31^ for transcription factors (Fig. 3A-D). This is likely because these two resources are manual efforts of extracting gene and protein sets from publications, including data from supporting tables. Comparing the kinase and transcription factor Rummagene libraries to PTMsigDB and ChEA 2022, Rummagene is likely more comprehensive and updated, but less accurate.

**Fig. 2.**
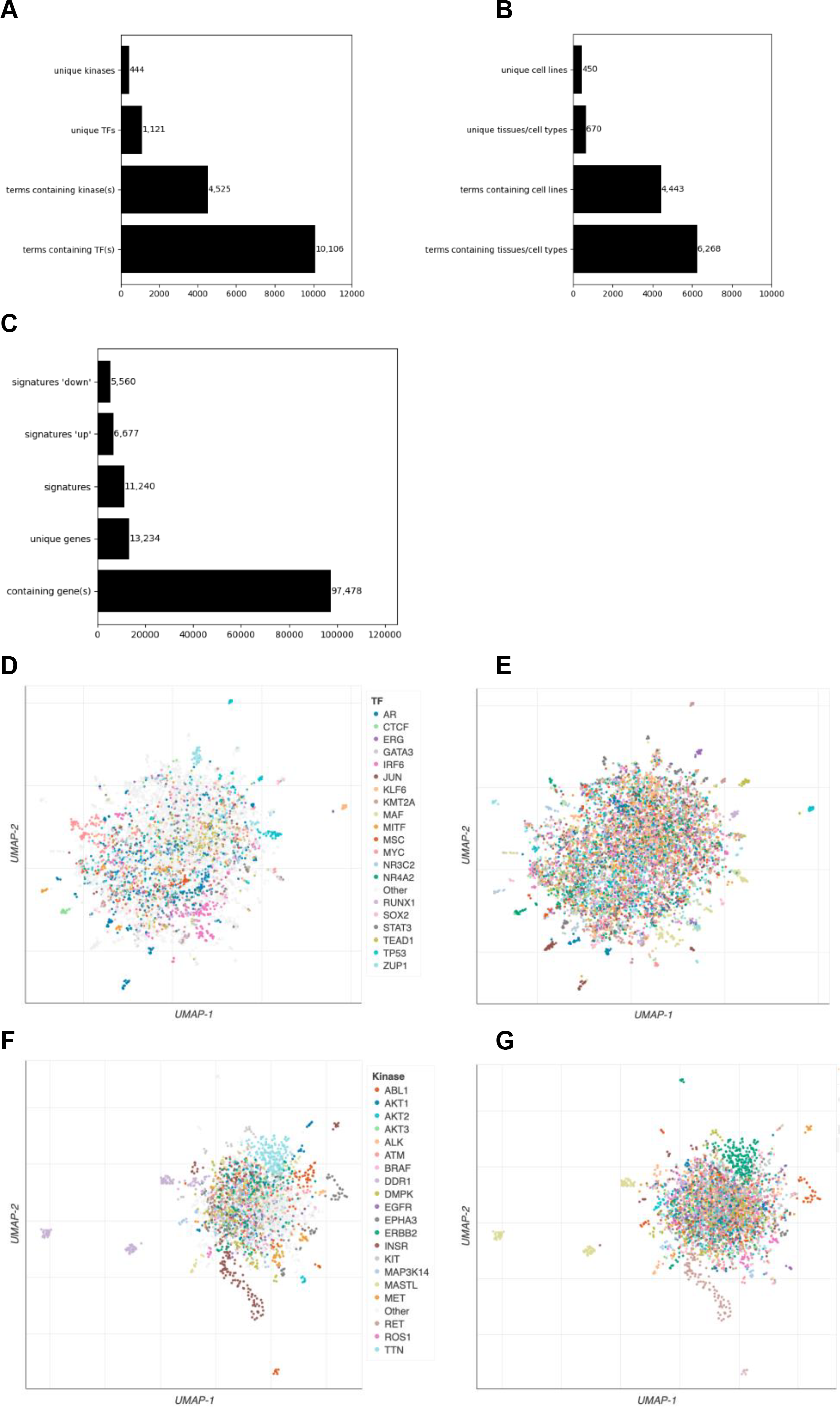
Extracting kinases, transcription factors, tissues, cell types, and cell lines from Rummagene gene sets. **A**. The number of unique and non-unique kinases and transcription factors identified in table headers describing gene sets. **B**. The number of unique and non-unique cell types and tissues, as well as cell lines identified in table headers describing gene sets. **C**. Gene set table titles containing the terms “up”, “down”, or gene names. **D**. UMAP projection of the transcription factors gene set library created from Rummagene where sets are colored by the topmost common transcription factors. **E**. The same UMAP as D except that sets are colored by their PMCID. **F**. UMAP projection of the kinases gene set library created from Rummagene where sets are colored by the topmost common kinases. **G**. The same UMAP as F except that sets are colored by their PMCID.

**Fig. 3.**
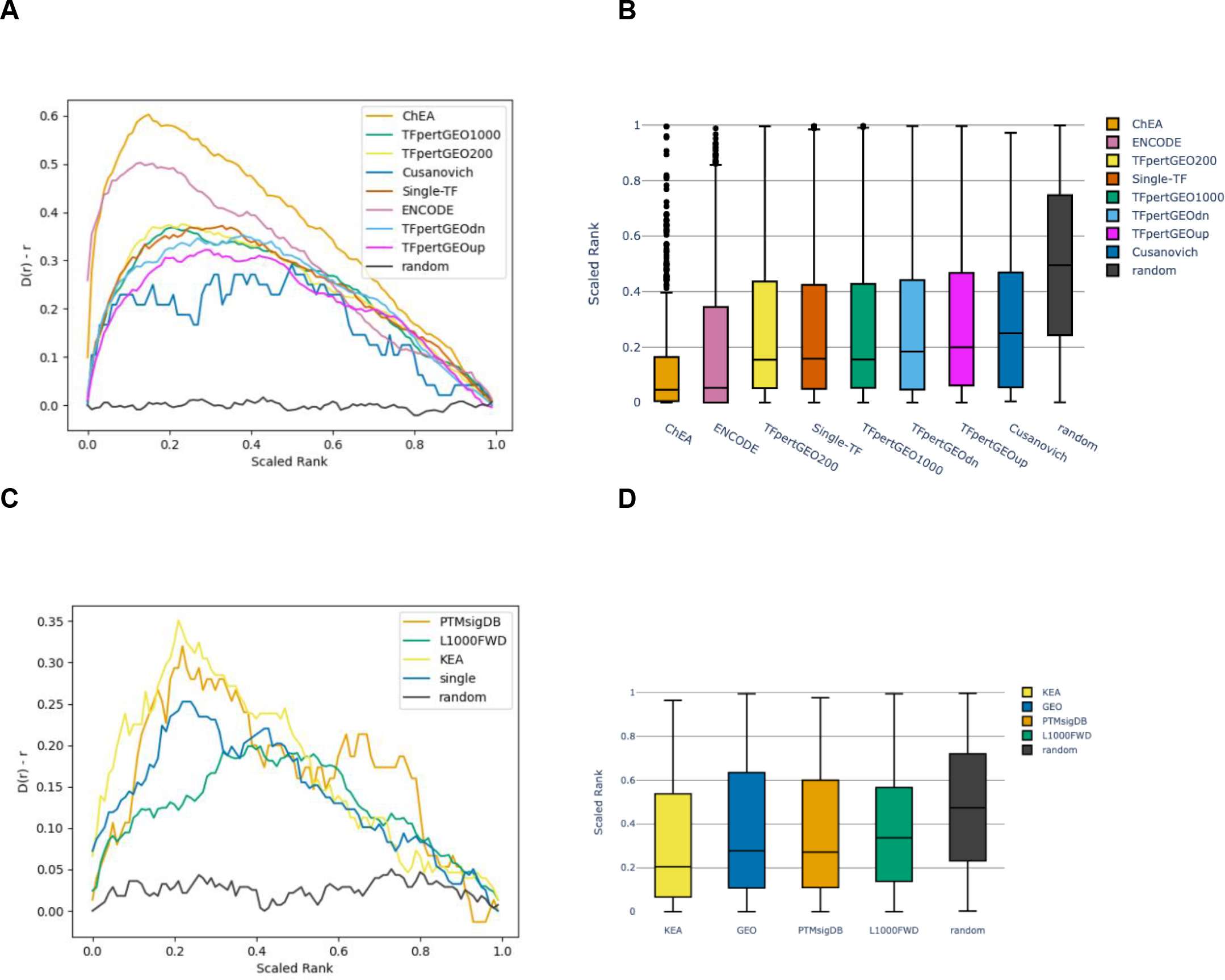
Benchmarking the consensus transcription factor gene and kinase set libraries created from Rummagene. **A**. The deviation of the cumulative distribution from uniform of the scaled ranks of each perturbed transcription factor in the benchmarking dataset. Kolmogorov-Smirnov test for goodness of fit for uniform distribution: ChEA (ChEA_2022) P = 1.04E-183; TFpertGEO1000 (TFpertGEO1000) P = 6.41E-113; TFpertGEO200 (TFpertGEO200) P = 1.35E-110; Cusanovich (Cusanovich_shRNA_TFs) P = 8.47E-06; Single-TF (Single-TF_Perturbations) P = 1.24E-118; ENCODE (ENCODE_TF_ChIP-seq_2015) P = 1.49E-128; TFpertGEOdn P = 1.24E-96; TFpertGEOup P = 6.28E-87; **B**. Scaled ranks of each transcription factor in the benchmarking datasets when enriched against the consensus TF gene set library. **C**. The deviation of the cumulative distribution from uniform of the scaled ranks of each perturbed kinase in the benchmarking datasets Kolmogorov-Smirnov test for goodness of fit for uniform distribution: PTMsigDB (PTMsigDB_drugtarget_signatures) P= 1.31E-04; L1000FWD (L1000FWD_kin_targets_updn) P = 2.19E-25; KEA (KEA_2015) P = 1.38E-14; GEO (single_kinase_perts_from_GEO_updn) P = 3.73E-15. **D**. Scaled ranks of each TF in the benchmarking datasets when enriched against the consensus kinase gene set library.

### Topic modeling

To obtain a global view of the contents of the gene sets in Rummagene, we performed Latent Dirichlet allocation (LDA) analysis^22^ on all abstracts from publications containing at least one extracted gene set. Nine topics were identified and subsequently manually labeled based on the most common terms and their relative weights (Fig. 4A). Some of the most frequently appearing terms across all topics included gene, cell, expression, DNA, patient, cancer, and analysis. The greatest portion of abstracts are relating to mutations and variants in diseases, protein-protein interactions, and mechanisms, while the topics with the least abstracts are related to immune functions and genome-wide associations and risks. The visualization of abstracts in topic space also reveals the relation and similarity between topics (Fig. 4B). For instance, the topic mutations and variants in disease borders DNA transcription and methylation. Additionally, the genome wide association and risk topic is isolated from the other topic clusters. The data and modeling topic is located adjacent to most of the other topics suggesting that abstracts with this topic may be related to a variety of other topics as expected. Overall, the topic analysis reveals the predominant categories of gene sets in Rummagene, specifically those concerning mutations and variants in diseases and those concerning protein interactions and functional mechanisms. Next, we explored abstracts that share high gene set similarity but have abstracts that are dissimilar.

**Fig. 4.**
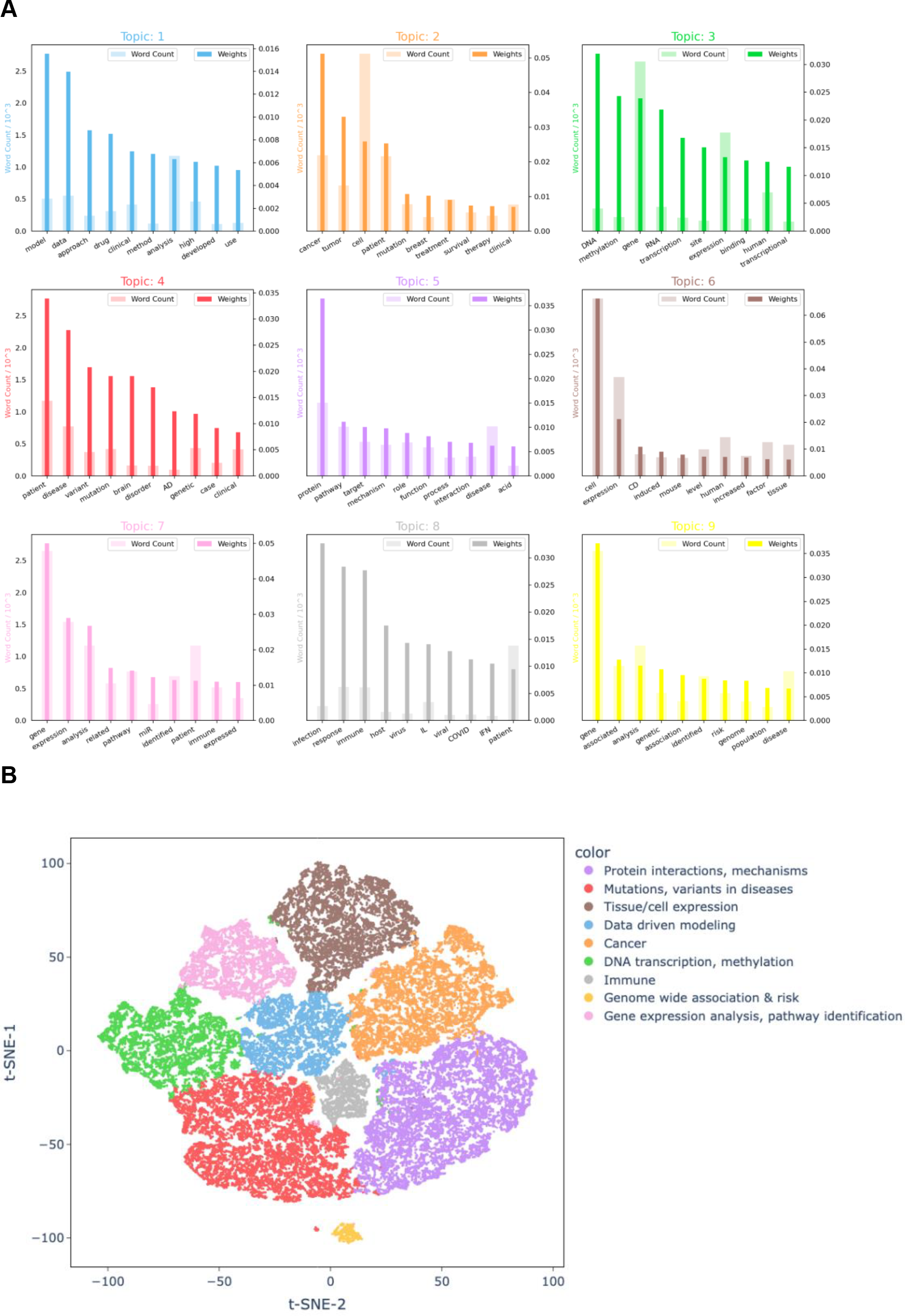
Topic modeling with Latent Dirichlet Allocation (LDA) from the abstracts of 121,043 papers from which gene sets in Rummagene were extracted. **A**. Based on the word counts and importance of each word from the LDA model, topics were manually labeled: Topic 1: Data driven modeling; Topic 2: Cancer; Topic 3: DNA transcription, methylation; Topic 4: Mutations, variants in diseases; Topic 5: Protein interactions, mechanisms; Topic 6: Tissue/cell expression; Topic 7: Gene expression analysis, pathway identification; Topic 8: Immune; Topic 9: Genome wide associations & risk. **B**. t-SNE projection of each of the 121,043 papers from which gene sets were extracted labeled by their topic identified through latent dirichlet allocation (LDA).

### Similar gene set pairs that are distant in abstract space

Next, we asked whether the knowledge embedded in Rummagene can lead to new hypotheses by identifying gene sets with high similarity in gene set space while completely disjointed at the publication abstract text space. The rationale for this is that this way we can identify undiscovered novel associations between named entities such as genes and diseases. Surprisingly, we first observed that the pairs of gene sets with the highest similarity at the gene set level, with no similarity at the abstract level, are highly enriched in proteins that are commonly detected in mass-spectrometry proteomics studies (Fig. 5A), highly expressed in RNA-seq assays (Fig. 5B), and widely studied (Fig. 5C). This is likely because proteomics studies tend to commonly report the same abundant, large-size, and “sticky” proteins, transcriptomics studies detected as differentially expressed highly expressed genes, and gene sets in publications commonly report overlapping genes in pathways and ontology terms containing highly studied genes. After filtering pairs of gene sets that are proteomics rich, or contain highly expressed genes, or composed of highly studied genes, we identified a few pairs of sets that contain a gene term in one table title of one set, and a disease term in the table title of the second set (Support text 1). For example, some of the top identified pairs highlight a possible relationship between the proteins identified to interact with CLUH^32^, and gene sets identified in hypoxia^33^, melanoma^34^, and glioma^35^. This connection is logical because CLUH was found to be critical to mitochondrial function which is altered in these conditions. Similarly, other top overlapping pairs include the TOPBP1 interactome^36^ and potential relationship to melanoma^34^, hypoxia^37^, and teratomas^38^. To assist in possibly explaining these connections, we utilized the GPT4 API, a large language model, to compose hypotheses that suggest how such seemingly unrelated named entities might be in fact related by giving GPT4 the two abstracts. The summaries produced by GPT4 are mostly helpful and logical but should be manually verified.

**Fig. 5.**
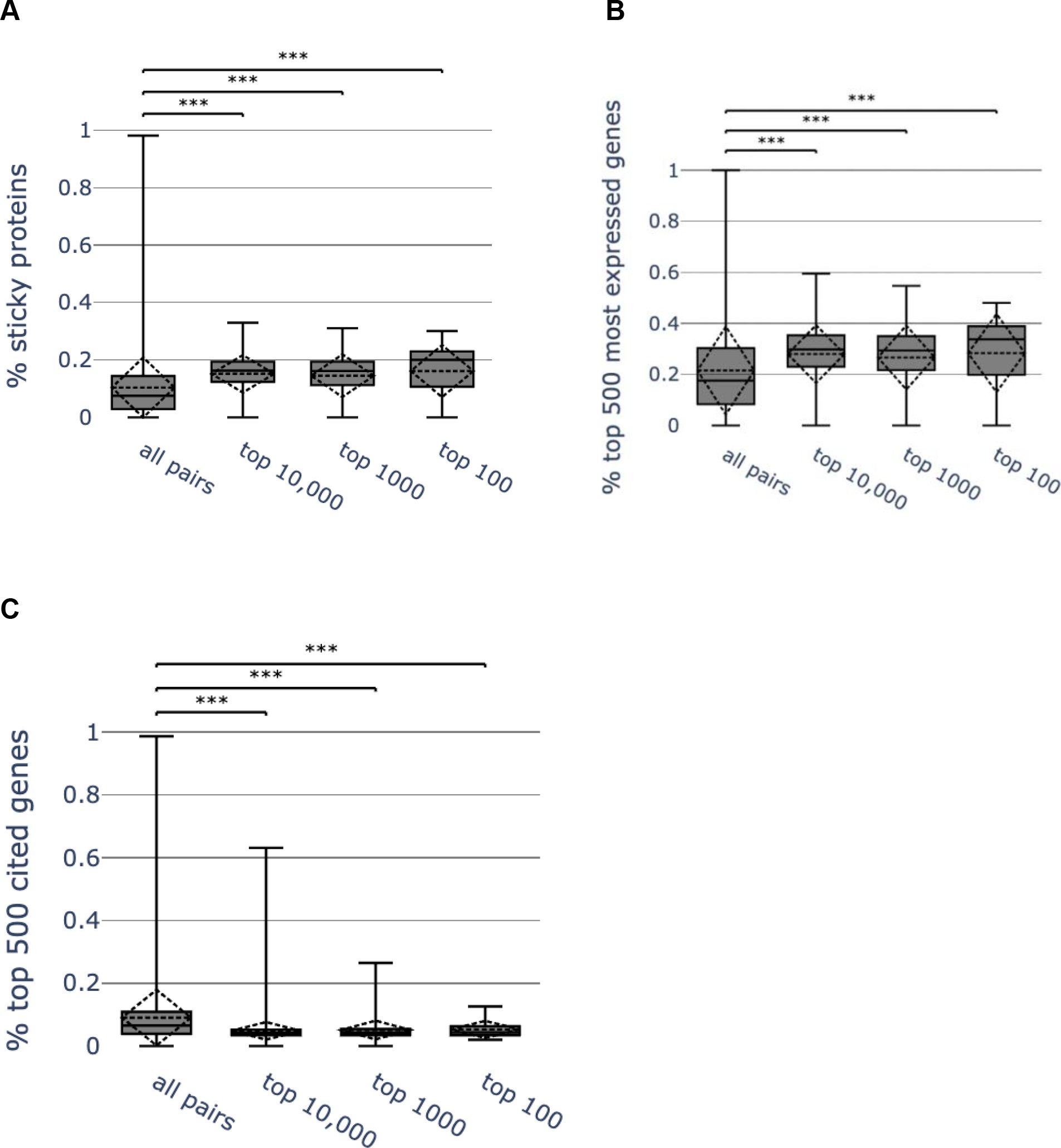
Distribution of percent of ‘sticky proteins, the percent top 500 most cited genes from GeneRIF, and the percent of top 500 most highly expressed gene sourced from ARCHS4, in the overlapping genes of the gene set pairs. *** indicates P values < 0.001. P values computed using the Welch’s t-test: **A**. % sticky protein: all pairs vs. top 10,000 P value = 0.00E+00; all pairs vs. top 1000 P value = 1.36E-60; all pairs vs. top 100 P value = 8.21E-09; **B**. % top 500 most expressed genes: all pairs vs. top 10,000 P value = 0.00E+00; all pairs vs. top 1000 P value = 3.73E-35; all pairs vs. top 100 P value = 2.44E-05; **C**. % top 500 most cited genes: all pairs vs. top 10,000 P value = 0.00E+00; all pairs vs. top 1000 P value = 5.87E-220; all pairs vs. top 100 P value = 1.02E-24.

### Gene function predictions

Large collections of gene sets can be used to effectively predict gene functions with semi-supervised learning^39^. The first step to produce such predictions is to construct a gene-gene similarity matrix from the Rummagene database of gene sets. This can be done with different algorithms. Here we tested the ability of three previously published co-occurrence algorithms^26^ to make such predictions, and compare the quality of such predictions to predictions made with a similar method that utilizes gene-gene co-expression correlations from thousands of RNA-seq samples^6^. The gene-gene similarity matrices from Rummagene were able to predict with high accuracy and precision gene membership for functional terms created from the Gene Ontology^40^, GWAS Catalog^41^, MGI MP^42^, and WikiPathways^43^ (Fig. 6A). To illustrate an example for one term, the term “Fasting Plasma Glucose” from GWAS Catalog was selected. The top 10 genes that are closest to the genes known to be associated with this phenotype are SLCO1B3-SLCO1B7, P3R3URF-PIK3R3, SLC30A8, FAM240B, MTNR1B, PERCC1, EEF1AKMT4-ECE2, KLF14, CCDC201, and PAX4; and the ROC curve to assess the quality of the predictions has a 0.75 area under the curve (Fig. 6B). The top 10 predicted genes for each term from these three gene set libraries are provided as a supporting table (Table S1).

**Fig. 6.**
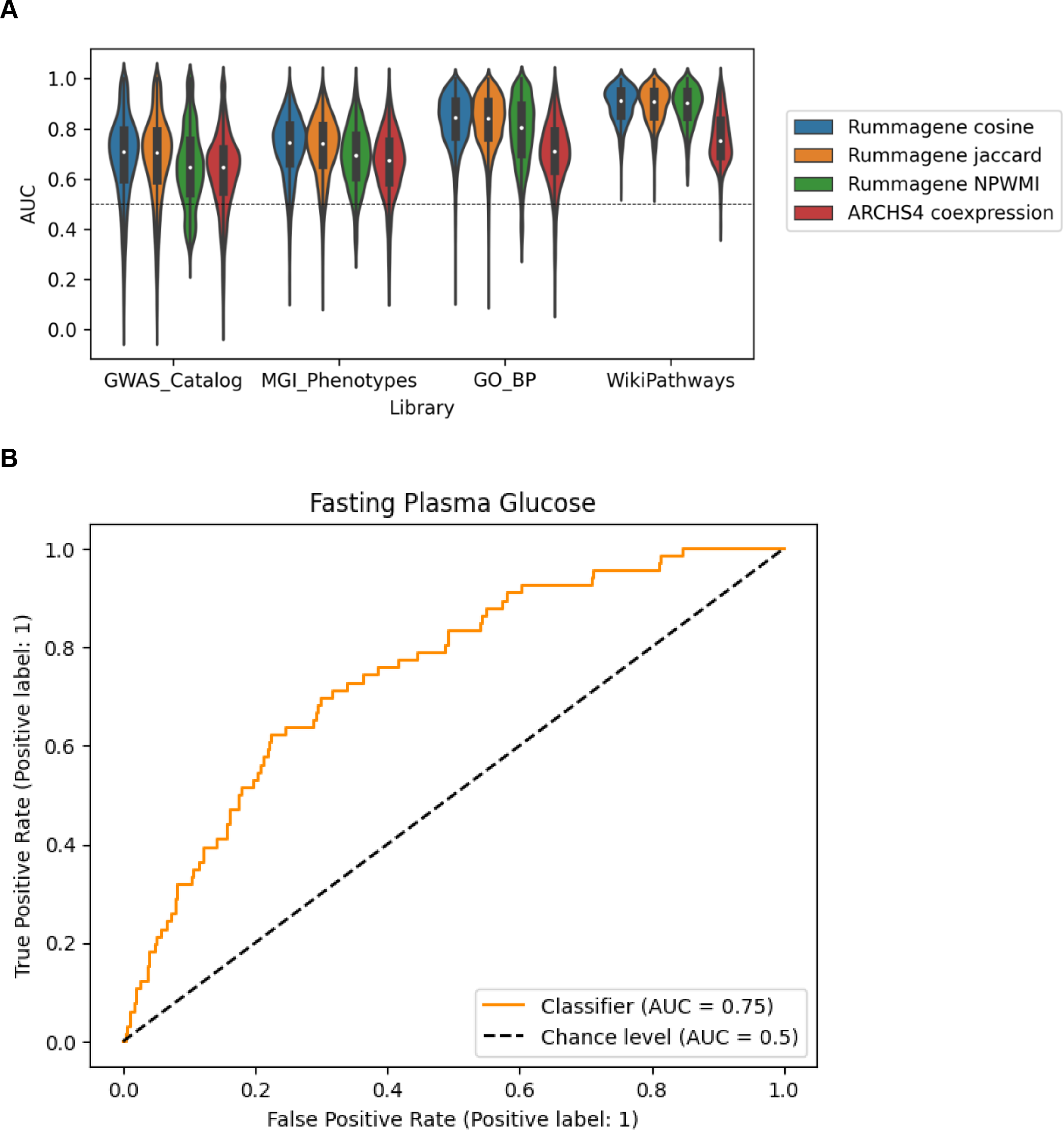
Benchmarking gene function prediction using Rummagene gene sets. **A**. The area under the receiver operating characteristic (AUROC) curve distributions for predicting genes associated with terms from four different gene-set libraries. Predictions were made using three gene-gene similarity matrices (cosine, Jaccard, and NPWMI) and the ARCHS4 gene-gene co-expression matrix. **B**. The AUROC curve for the GWAS Catalog term “Fasting Plasma Glucose”, produced from the average NPWMI of each gene to the 66 genes included in the “Fasting Plasma Glucose” gene set.

### The knowledge space that is covered by Rummagene compared with Enrichr

To assess the breadth and coverage of the automatically curated Rummagene gene set space, we contrasted it against the Enrichr^31^ gene set space. Enrichr is a large-scale curated database of gene sets of similar size when compared to Rummagene. UMAP^23^ was applied to project over 1 million gene sets into two dimensions for the purpose of data visualization where each point represents a gene set from either Rummagene or Enrichr. Gene sets are colored by whether they originate from Rummagene or Enrichr’s gene set library categorization: Transcription, Pathways, Ontologies, Diseases/Drugs, Cell Types, Misc, Legacy, Crowd (Fig. 7A-C). We observe that Rummagene gene sets cluster into many punctate clusters that likely represent themed gene sets (Fig. 7A). Also, Enrichr’s gene sets are clustered by category (Fig. 7B). When overlaying the Rummagene gene sets on the Enrichr gene set, most categories are covered with some few exceptions. We observe that the gene set libraries created from the LINCS L1000 data^44^ and from DrugMatrix^45^ are not covered by Rummagene, while few areas in gene set space are much more common in Rummagene compared with Enrichr.

**Fig. 7.**
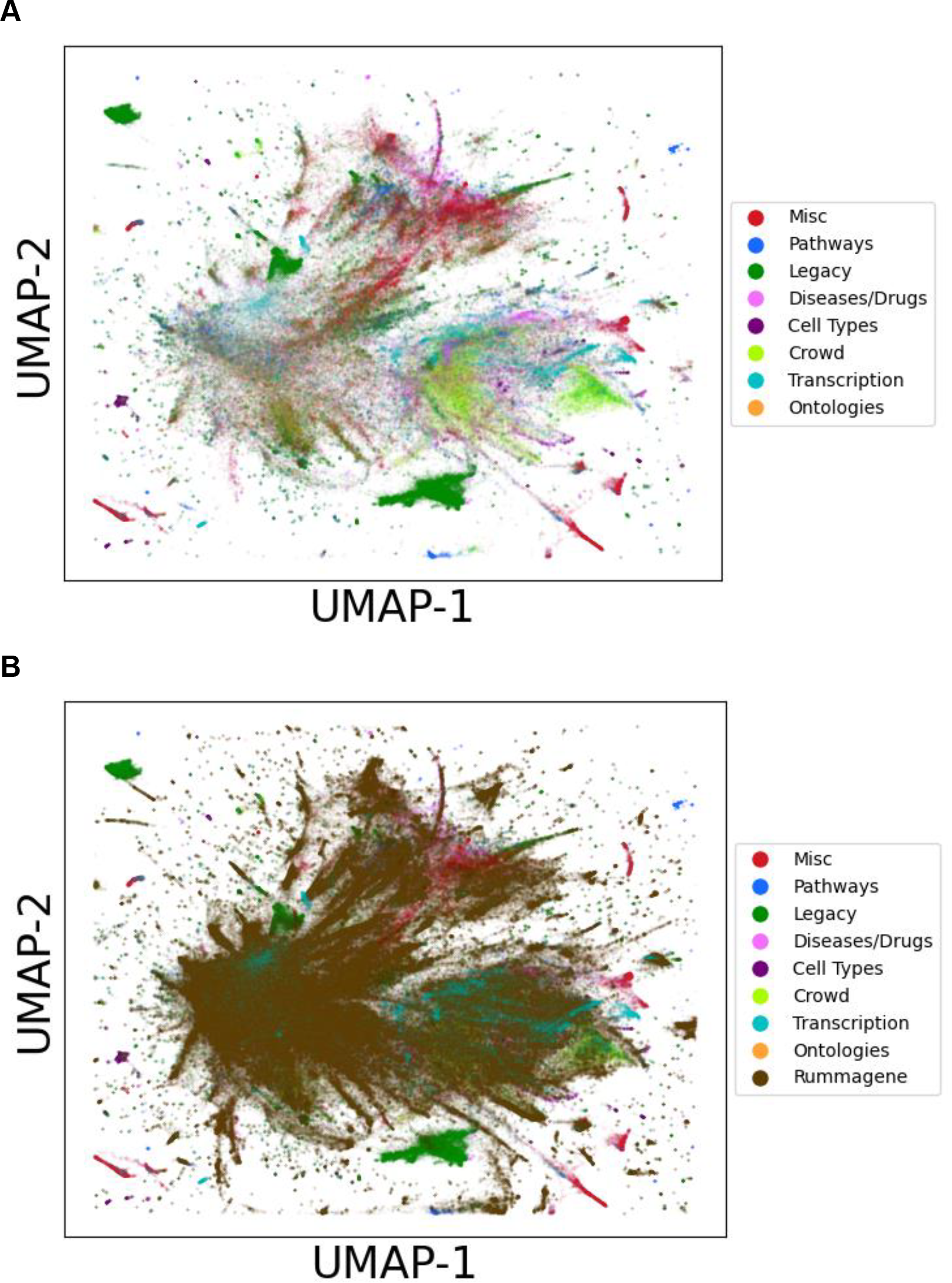
Visualizing the global space of the gene sets contained within Rummagene and Enrichr. Vectorized by IDF followed by Truncated SVD to 50 dimensions is applied to the combined gene sets from both Rummagene and Enrichr. The Enrichr gene sets are colored by library categories. UMAP visualization of the gene sets in Enrichr without **A**. and with **B**. the Rummagene sets. 10% of extreme points are omitted for zooming into the area with the most sets/points.

### The Rummagene website

The Rummage data is served on the website https://rummagene.com with three search engines. The first search engine accepts gene sets as the input query. This search engine returns matching gene sets based on the overlap between the input gene set and the sets in the Rummagene database. The results are ranked by the Fisher exact test, and to optimize responsiveness, a fast in-memory algorithm is implemented. The results are presented to the user in paginated tables with hyperlinks to the original publications from which gene sets were extracted, the genes in the matching sets, the overlapping genes, the p-values and corrected p-values of the overlap, and the odds ratios. The numbers of overlapping genes and genes in the matching sets are also hyperlinked. When clicking these numbers, a popup screen shows the genes with the ability to copy them to the clipboard, submit them to Rummagene, or submit them to Enrichr^31^. The second search engine facilitates a free-text PMC search. This search engine queries PMC with the entered terms to receive PMCIDs that match the query. It then compares the returned PMCIDs to the PMCIDs in Rummagene to identify matching PMCIDs. Once such matches are detected, the gene sets in the Rummagene database are returned to the user as a paginated table with hyperlinks to the original publications and the matching gene sets. The third search engine queries the table titles from which gene sets were extracted. Table titles that match the inputted search terms are displayed in a paginated table with hyperlinks to the matching publications and gene sets. The entire database is available for download as a text file, and access to the data is provided via API. Importantly, the Rummagene resource is updated automatically once a week.

## Discussion and Conclusions

By crawling through full articles and supporting materials from over five million research publications available from PMC, we were able to identify over 150,000 publications that contain over 600,000 mammalian gene sets. Such collections of gene sets contain gene sets of various lengths. Smaller gene sets are enriched for widely studied genes while longer lists contain less studied genes likely due to their origin from omics studies. Interesting, in the past five years, the publication of gene sets in articles has been increasing exponentially. Hence, most gene sets in the Rummagene database are from this period. Here we demonstrated how the Rummagene resource can be used for various applications. Specifically, we showed how a subset of the extracted sets can be used for transcription factor, kinase, cell type and tissue, cell line, and signature enrichment analyses. We also showed how the rich knowledge in Rummagene can be used for gene function predictions. In addition, we demonstrated how we can form novel hypotheses by identifying gene set pairs with high similarity in gene set space and low similarity in abstract space. However, many additional applications are possible. For example, Rummagene can be used for universal scRNA-seq cell type identification, or for producing textual descriptions for gene sets using large language models (LLMs). One of the opportunities provided by Rummagene is its integration with other resources that contain large collections of gene sets and signatures, for example, Enrichr^31^, ARCHS4^6^, and SigCom LINCS^46^. Biomedical research has been traditionally communicated via hardcopy printed paper journals. The transition into fully digital research communication, and with the introduction of omics technologies, increased efforts are placed on better annotation and standardization of published research data including the publication of gene sets and data tables. During this transition period toward such improved annotations, Rummagene plays a significant role in making previously published data, buried in supplemental materials of publications, more findable, accessible, interoperable, and reusable (FAIR)^47^.

## Acknowledgements

This study is partially supported by NIH grants OT2OD030160, U24CA264250, RC2DK131995, and U24CA224260.

