## Supporting Text 1 for "Rummagene: Mining Gene Sets from Supporting Materials of PMC Publications"

1. PMC7981264-NIHMS1666748-supplement-1666748_Supp_Data6.xlsx-Glioma_grade-Unnamed_2,
PMC8744257-12915_2021_1213_MOESM7_ESM.xls-HCT116_BioID_CLUH-Unnamed_35

p-value: 1.9602455624474105e-306, odds ratio: 7.481400437636761, overlap: 897, percent sticky protein: 0.0869565217391304
GS2: CLUH, GS1: glioma
0 publications identified co-mentioning the terms:
https://pubmed.ncbi.nlm.nih.gov/?term=CLUH+glioma&sort=date

**Hypothesis: The high overlap between the two gene sets, despite being extracted from two publications with dissimilar abstracts, may be due to the role of the CLUH gene in mitochondrial function and its potential influence on the progression of glioma.**

The first paper discusses the proteomic signatures associated with more aggressive cancers, including glioma. It mentions that these signatures involve DNA copy number alterations and pathways of altered metabolism, Warburg-like effects, and translation factors. These factors could potentially be influenced by the function of mitochondria, which are essential for energy production and other fundamental biological processes.

The second paper focuses on the CLUH gene, an RNA binding protein (RBP) that specifically recognizes mRNAs coding for mitochondrial proteins. The paper reveals that CLUH interacts with mitochondrial proteins and their cognate mRNAs in the cytosol during the process of active translation. This suggests that CLUH may play a crucial role in the regulation of mitochondrial function.

Given the importance of mitochondria in cancer progression, it is plausible that the CLUH gene could influence the progression of glioma by affecting mitochondrial function. This could explain the high overlap between the two gene sets. Further research would be needed to confirm this hypothesis and to explore the specific mechanisms through which CLUH may influence glioma progression.

2. PMC8896345-Table_1.xlsx-SKOV3_Hypoxia_Phosphopeptides-Gene,
PMC8744257-12915_2021_1213_MOESM7_ESM.xls-HCT116_BioID_CLUH-Unnamed_35

p-value: 1.9544511337218995e-281, odds ratio: 12.368003412011626, overlap: 576, percent sticky protein: 0.0729166666666666
GS2: CLUH, GS1: hypoxia
0 publications identified co-mentioning the terms:
https://pubmed.ncbi.nlm.nih.gov/?term=CLUH+hypoxia&sort=date

**Hypothesis: The high overlap between the two gene sets, despite being extracted from two publications with dissimilar abstracts, may be due to the potential role of the CLUH gene in hypoxia adaptation in ovarian cancer cells.**

The first paper discusses the adaptive potential of ovarian cancer cells, specifically SKOV-3 cells, under hypoxic conditions. Hypoxia, or low oxygen levels, is a common feature in solid tumors like ovarian cancer. The study highlights the role of post-translational protein modifications in the adaptive response of these cells to hypoxia.

The second paper focuses on the CLUH gene, an RNA binding protein (RBP) that specifically recognizes mRNAs coding for mitochondrial proteins. Mitochondria are essential for energy production in cells and their function can be significantly affected under hypoxic conditions. The paper suggests that CLUH may be involved in the regulation of mitochondrial protein translation and stability, which could be crucial for cell survival and adaptation under stress conditions such as hypoxia.

Therefore, it is plausible that the CLUH gene may be involved in the adaptive response of SKOV-3 ovarian cancer cells to hypoxia, possibly by regulating the translation and stability of mitochondrial proteins. This could explain the high overlap between the two gene sets. Further experimental studies would be needed to confirm this hypothesis.

3. PMC8111777-mmc2.xlsx-V_melanoma-Unnamed_6,
PMC8744257-12915_2021_1213_MOESM7_ESM.xls-HCT116_BioID_CLUH-Unnamed_35

p-value: 3.1792579226647546e-251, odds ratio: 7.494487179487179, overlap: 707, percent sticky protein: 0.0792079207920792
GS2: CLUH, GS1: melanoma
0 publications identified co-mentioning the terms:
https://pubmed.ncbi.nlm.nih.gov/?term=CLUH+melanoma&sort=date

**Hypothesis: The high overlap between the two gene sets, despite being extracted from publications with dissimilar abstracts, may be due to the role of the CLUH gene in the regulation of mitochondrial proteins and the potential involvement of these proteins in melanoma progression.**

The first abstract discusses a study that uses AGPC-based protein extraction to profile phosphorylations in the DNA damage response pathway after ionizing irradiation of U2OS cells. This method was found to be effective for obtaining high-quality phosphosite data from cells and tissue samples, including tumor tissues. Given that melanoma is a type of tumor, it is plausible that the gene set extracted from this study includes genes involved in the DNA damage response pathway, which may be relevant to melanoma progression.

The second abstract focuses on the CLUH gene, an RNA binding protein (RBP) that specifically recognizes mRNAs coding for mitochondrial proteins. The study reveals that CLUH interacts with these mRNAs during the process of active translation, suggesting a role for CLUH in the regulation of mitochondrial protein synthesis. Given the central role of mitochondria in energy production and other fundamental biological processes, it is possible that alterations in mitochondrial protein synthesis could contribute to disease processes, including melanoma.

Therefore, the high overlap between the two gene sets may reflect the involvement of the CLUH gene and its target mitochondrial proteins in melanoma progression. Further research is needed to confirm this hypothesis and elucidate the precise mechanisms underlying this potential connection.

4. PMC9270554-12672_2022_524_MOESM7_ESM.xlsx-Top2000DEG-Supplemental_Table_1_-_Top_2000_differentially_expressed_genes_in_BC-K562_cells_under_hypoxia,
PMC6949271-41467_2019_13981_MOESM4_ESM.xlsx-TOPBP1-TOPBP1_interacting_proteins_identified_by_MS_MS

p-value: 3.1942264381642472e-239, odds ratio: 6.510819118547449, overlap: 758, percent sticky protein: 0.0870712401055409
GS2: MS+TOPBP1, GS1: hypoxia
0 publications identified co-mentioning the terms:
https://pubmed.ncbi.nlm.nih.gov/?term=MS+TOPBP1+hypoxia&sort=date

**Hypothesis: The high overlap between the two gene sets, despite being extracted from two publications with dissimilar abstracts, could be due to the potential role of the TOPBP1 gene in the cellular response to hypoxic conditions, which is the disease term from the first gene set.**

The first paper discusses the impact of hypoxia on metabolic reprogramming and plasticity in chronic myeloid leukaemia (CML) cells. Hypoxia, or low oxygen conditions, is known to induce various cellular responses, including changes in gene expression. The paper specifically mentions the differential expression of genes (DEG) under hypoxia related to various cellular processes such as the Krebs cycle, lipid synthesis, cholesterol homeostasis, mitophagy, and mitochondrial biogenesis.

The second paper discusses the role of the TOPBP1 gene in the cellular response to DNA double-strand breaks (DSBs) in ribosomal DNA (rDNA) repeats. TOPBP1 is recruited to the nucleoli in response to rDNA breaks, leading to inhibition of ribosomal RNA synthesis and nucleolar segregation.

Given these findings, it is plausible that the TOPBP1 gene could be involved in the cellular response to hypoxia. Hypoxia can induce DNA damage, including DSBs, due to the increased production of reactive oxygen species (ROS). Therefore, the TOPBP1 gene, which is involved in the cellular response to DSBs, could be differentially expressed under hypoxic conditions, leading to its inclusion in the gene set from the first paper.

Furthermore, the inhibition of ribosomal RNA synthesis and nucleolar segregation, which are regulated by TOPBP1, could be part of the cellular response to hypoxia, contributing to the metabolic reprogramming and plasticity observed in the CML cells.

Therefore, despite the dissimilar abstracts, the potential role of the TOPBP1 gene in the cellular response to hypoxia could explain the high overlap between the two gene sets. However, further research is needed to confirm this hypothesis and elucidate the exact mechanisms involved.

5. PMC9270554-12672_2022_524_MOESM7_ESM.xlsx-Top2000DEG-Supplemental_Table_1_-_Top_2000_differentially_expressed_genes_in_BC-K562_cells_under_hypoxia,
PMC6949271-41467_2019_13981_MOESM4_ESM.xlsx-TOPBP1-Unnamed_3

p-value: 3.1942264381642472e-239, odds ratio: 6.510819118547449, overlap: 758, percent sticky protein: 0.0870712401055409
GS2: TOPBP1, GS1: hypoxia
1 publications identified co-mentioning the terms:
https://pubmed.ncbi.nlm.nih.gov/?term=TOPBP1+hypoxia&sort=date

**Hypothesis: The high overlap between the two gene sets, despite being extracted from two publications with dissimilar abstracts, could be due to the role of TOPBP1 gene in the cellular response to hypoxia, a condition mentioned in the first abstract.**

The first abstract discusses the effects of hypoxia on chronic myeloid leukaemia (CML) cells, specifically BC-K562 cells. Hypoxia, a condition characterized by low oxygen levels, can trigger metabolic reprogramming and plasticity in cells, affecting their survival and response to certain drugs. The abstract mentions that hypoxia drives differential gene expression related to various cellular processes such as the Krebs cycle, lipid synthesis, cholesterol homeostasis, mitophagy, and mitochondrial biogenesis.

The second abstract discusses the role of the TOPBP1 gene in the cellular response to DNA double-strand breaks (DSBs) in ribosomal DNA (rDNA) repeats. TOPBP1 is recruited to the nucleoli in response to rDNA breaks, and this recruitment is necessary for the inhibition of ribosomal RNA synthesis and nucleolar segregation.

The connection between the two gene sets could be that the TOPBP1 gene plays a role in the cellular response to hypoxia. Hypoxia can cause DNA damage, including DSBs, and the cellular response to this damage could involve the recruitment of TOPBP1 to the nucleoli. This could result in changes in gene expression, including the differential expression of genes related to the cellular processes mentioned in the first abstract. Therefore, despite the dissimilar abstracts, the two gene sets could have a high overlap due to the shared involvement of the TOPBP1 gene in the cellular response to hypoxia.

6. PMC6075738-NIHMS977848-supplement-6.xlsx-Teratomas_to_Pure_Embryonals-Unnamed_11,
PMC6949271-41467_2019_13981_MOESM4_ESM.xlsx-TOPBP1-TOPBP1_interacting_proteins_identified_by_MS_MS

p-value: 1.723406896277315e-183, odds ratio: 4.910391191313656, overlap: 747, percent sticky protein: 0.0963855421686747
GS2: MS+TOPBP1, GS1: teratoma
0 publications identified co-mentioning the terms:
https://pubmed.ncbi.nlm.nih.gov/?term=MS+TOPBP1+teratoma&sort=date

**Hypothesis: The high overlap between the two gene sets, despite being extracted from two publications with dissimilar abstracts, could be due to the potential role of the TOPBP1 gene in the development or progression of teratomas.**

The first abstract discusses the molecular characteristics of testicular germ cell tumors (TGCTs), including teratomas. It mentions the significance of somatic mutations in three genes, including KIT, KRAS, and NRAS, in samples with seminoma components. The abstract also highlights the role of epigenomic processes in determining histologic fates in TGCTs, suggesting that gene expression and regulation play a crucial role in the development of these tumors.

The second abstract focuses on the role of the TOPBP1 gene in the response to DNA double-strand breaks (DSBs) in ribosomal DNA (rDNA) repeats. It suggests that the recruitment of TOPBP1 in the nucleoli is required for inhibition of ribosomal RNA synthesis and nucleolar segregation in response to rDNA breaks.

Given the role of TOPBP1 in DNA repair and the importance of gene mutations in the development of teratomas, it is plausible that mutations or dysregulation of TOPBP1 could contribute to the development or progression of teratomas. This could explain the high overlap between the two gene sets. Further research would be needed to confirm this hypothesis and elucidate the exact mechanisms involved.

7. PMC6075738-NIHMS977848-supplement-6.xlsx-Teratomas_to_Pure_Embryonals-Unnamed_11,
PMC6949271-41467_2019_13981_MOESM4_ESM.xlsx-TOPBP1-Unnamed_3

p-value: 1.723406896277315e-183, odds ratio: 4.910391191313656, overlap: 747, percent sticky protein: 0.0963855421686747
GS2: TOPBP1, GS1: teratoma
0 publications identified co-mentioning the terms:
https://pubmed.ncbi.nlm.nih.gov/?term=TOPBP1+teratoma&sort=date

**Hypothesis: The high overlap between the two gene sets, despite being extracted from two publications with dissimilar abstracts, may be due to the potential role of the TOPBP1 gene in the development or progression of teratomas.**

The first paper discusses the molecular characteristics of different types of testicular germ cell tumors (TGCTs), including teratomas. It mentions the significant somatic mutation of three genes—KIT, KRAS, and NRAS—in samples with seminoma components. However, the paper does not mention the TOPBP1 gene directly. It does, however, highlight the role of epigenomic processes in determining histologic fates in TGCTs, suggesting that various genes and their interactions may be involved in the development of these tumors.

The second paper focuses on the role of the TOPBP1 gene in the response to DNA double-strand breaks (DSBs) in ribosomal DNA (rDNA) repeats. It suggests that the recruitment of TOPBP1 in the nucleoli is required for inhibition of ribosomal RNA synthesis and nucleolar segregation in response to rDNA breaks. This indicates that TOPBP1 plays a crucial role in the cellular response to DNA damage.

Given the role of TOPBP1 in DNA damage response, it is plausible that alterations in this gene could contribute to the development or progression of teratomas. DNA damage and improper repair mechanisms are known to be involved in tumorigenesis. Therefore, it is possible that the TOPBP1 gene could be a common factor in the gene sets from both papers, linking the disease term "teratoma" with the gene term "TOPBP1". Further research would be needed to confirm this hypothesis and elucidate the exact role of TOPBP1 in teratomas.

8. PMC4490555-ncomms8289-s3.xls-Dynamic_Heat_Shock_Targets-Gene_names, PMC8744257-12915_2021_1213_MOESM7_ESM.xls-HCT116_BioID_CLUH-Unnamed_35

p-value: 1.5670973587819978e-145, odds ratio: 9.469690440278674, overlap: 334, percent sticky protein: 0.059880239520958
GS2: CLUH, GS1: shock
1 publications identified co-mentioning the terms:
https://pubmed.ncbi.nlm.nih.gov/?term=CLUH+shock&sort=date

**Hypothesis: The high overlap between the two gene sets, despite being extracted from two publications with dissimilar abstracts, may be due to the role of the CLUH gene in the regulation of mitochondrial proteins and the response of cells to heat shock, which is a form of stress.**

In the first paper, the focus is on SUMOylation, a post-translational modification (PTM) that regulates all nuclear processes. The paper specifically discusses the identification of SUMOylation sites in response to heat shock. Heat shock is a form of stress that can cause significant changes in the cellular environment, including the alteration of protein function and structure. The term "shock" in this context is related to a sudden, stressful change in the environment that the cell has to respond to.

The second paper discusses the role of the CLUH gene, an RNA binding protein (RBP) that specifically recognizes mRNAs coding for mitochondrial proteins. The CLUH gene is involved in the regulation of mRNA translation, stability, or localization, which are all crucial processes for the proper functioning of mitochondria.

Given these findings, it's plausible that the CLUH gene may play a role in the cellular response to heat shock. Under stress conditions such as heat shock, cells need to rapidly adjust their protein synthesis machinery to ensure survival. As an RBP that regulates mRNAs coding for mitochondrial proteins, CLUH could be involved in this response by modulating the translation of these proteins.

Therefore, the high overlap between the two gene sets could be due to the involvement of the CLUH gene in the cellular response to heat shock, which could involve changes in SUMOylation patterns. Further investigation would be needed to confirm this hypothesis and elucidate the exact mechanisms involved.

9. PMC10293682-Table1.XLSX-S2-Supplementary_Table_2_RNA-binding_proteins_genomic_alterations_in_Colon_adenocarcinoma_TCGA_PanCancer_Atlas_and_CPTAC-2_Prospective,
PMC4691244-mmc2.xlsx-Sheet1-T7-DGCR8

p-value: 4.075141520104804e-142, odds ratio: 40.27695405454878, overlap: 166, percent sticky protein: 0.036144578313253
GS2: DGCR8, GS1: adenocarcinoma
24 publications identified co-mentioning the terms:
https://pubmed.ncbi.nlm.nih.gov/?term=DGCR8+adenocarcinoma&sort=date

**Hypothesis: The high overlap between the two gene sets, despite being extracted from two publications with dissimilar abstracts, could be due to the role of DGCR8 in RNA processing and its potential involvement in adenocarcinoma progression.**

The first gene set is related to RNA-binding proteins and their genomic alterations in colon adenocarcinoma. The abstract discusses the role of RNA-binding proteins (RBPs) in the progression of colon and rectal cancer, and the need to identify sensitive biomarkers for these cancers. It also mentions the identification of new RBPs involved in the progression of these cancers.

The second gene set is related to DGCR8, a gene involved in the biogenesis of microRNA (miRNA) and the regulation of the stability of several types of cellular RNAs. The abstract discusses the role of DGCR8 in the recruitment of the exosome to structured RNAs and the control of their stability.

The connection between the disease (adenocarcinoma) and the gene (DGCR8) could be that DGCR8, through its role in RNA processing, may be involved in the regulation of the RBPs identified in the first paper. These RBPs could be potential biomarkers for adenocarcinoma, and their regulation by DGCR8 could influence the progression of the disease. This connection could explain the high overlap between the two gene sets.

Further research is needed to validate this hypothesis and to elucidate the molecular mechanisms underlying the potential involvement of DGCR8 in adenocarcinoma progression.

10. PMC10293285-41467_2023_39210_MOESM9_ESM.xlsx-neuroblastoma_cell-Pathway, PMC7046753-Table_4.xlsx-Base_TF_FOS-TF_name_A

p-value: 7.442389652442631e-109, odds ratio: 44.31298681009663, overlap: 102, percent sticky protein: 0.0098039215686274
GS2: FOS, GS1: neuroblastoma
178 publications identified co-mentioning the terms:
https://pubmed.ncbi.nlm.nih.gov/?term=FOS+neuroblastoma&sort=date

**Hypothesis: The FOS gene, which is involved in transcription regulation, may play a significant role in the metastasis of neuroblastoma, contributing to the high overlap of the two gene sets despite the dissimilar abstracts.**

The first paper focuses on the metastasis of neuroblastoma, particularly to the bone marrow. It discusses the cellular plasticity of neuroblastoma tumor cells and their interaction with the bone marrow microenvironment. The paper also mentions the activation of pro- and anti-inflammatory programs and the expression of tumor-promoting factors, which are likely to involve various genes and pathways.

The second paper, on the other hand, discusses the use of Chromatin immunoprecipitation followed by next-generation sequencing (ChIP-Seq) to study regulatory DNA-protein interactions at the genetic and epigenetic level. It specifically mentions the FOS gene, which is known to be a transcription factor. Transcription factors are proteins that control the rate of transcription of genetic information from DNA to messenger RNA, by binding to a specific DNA sequence.

Given the role of the FOS gene in transcription regulation, it is plausible that this gene could be involved in the processes discussed in the first paper. For instance, the FOS gene could be involved in the regulation of genes that promote the metastasis of neuroblastoma cells, their cellular plasticity, or their interaction with the bone marrow microenvironment. This could explain the high overlap between the two gene sets, despite the dissimilar abstracts. Further research would be needed to confirm this hypothesis and elucidate the exact role of the FOS gene in neuroblastoma metastasis.

11. PMC8111777-mmc2.xlsx-V_melanoma-Unnamed_6, PMC6949271-41467_2019_13981_MOESM4_ESM.xlsx-TOPBP1-TOPBP1_interacting_proteins_identified_by_MS_MS

p-value: 6.3773377177697545e-102, odds ratio: 3.845804174013776, overlap: 510, percent sticky protein: 0.0980392156862745
GS2: MS+TOPBP1, GS1: melanoma
0 publications identified co-mentioning the terms:
https://pubmed.ncbi.nlm.nih.gov/?term=MS+TOPBP1+melanoma&sort=date

**Hypothesis: The high overlap between the two gene sets, despite being extracted from two publications with dissimilar abstracts, may be due to the involvement of both gene sets in the DNA damage response pathway, which is crucial in the development of diseases such as melanoma.**

The first paper discusses the use of acid guanidinium thiocyanate–phenol–chloroform (AGPC) for mass spectrometry–based phosphoproteomics, which is used to profile phosphorylations in the DNA damage response pathway. This is important in the context of melanoma, as disruptions in the DNA damage response pathway can lead to uncontrolled cell growth and cancer development. The paper specifically mentions the involvement of ATM, ATR, CHEK1/2, and PRKDC in this pathway, which are all key players in the DNA damage response.

The second paper focuses on the role of TOPBP1 and its interaction with other proteins in response to DNA double-strand breaks (DSBs) in ribosomal DNA (rDNA) repeats. TOPBP1 is known to play a crucial role in the DNA damage response, and its recruitment is dependent on both ATM and ATR activity. This suggests a potential link between the gene TOPBP1 and the disease melanoma, as both are associated with the DNA damage response pathway.

Therefore, despite the dissimilar abstracts, the high overlap between the two gene sets could be due to their shared involvement in the DNA damage response pathway. Further research could investigate the specific role of TOPBP1 in melanoma development and whether it could be a potential target for treatment.

12. PMC8111777-mmc2.xlsx-V_melanoma-Unnamed_6, PMC6949271-41467_2019_13981_MOESM4_ESM.xlsx-TOPBP1-Unnamed_3

p-value: 2.4521174914114427e-101, odds ratio: 3.832699781585952, overlap: 509, percent sticky protein: 0.0982318271119842
GS2: TOPBP1, GS1: melanoma
1 publication identified co-mentioning the terms:
https://pubmed.ncbi.nlm.nih.gov/?term=TOPBP1+melanoma&sort=date

**Hypothesis: The high overlap between the two gene sets, despite being extracted from two publications with dissimilar abstracts, may be due to the role of the TOPBP1 gene in the DNA damage response pathway, which is implicated in the development of melanoma.**

The first paper discusses a method for extracting and analyzing biomolecules, including DNA, RNA, and proteins, from small samples using acid guanidinium thiocyanate–phenol–chloroform (AGPC). This method was used to profile phosphorylations in the DNA damage response pathway after ionizing irradiation of U2OS cells. The DNA damage response pathway is crucial for maintaining genomic stability, and dysregulation of this pathway can lead to the development of diseases such as cancer, including melanoma.

The second paper focuses on the role of the TOPBP1 gene in the response to DNA double-strand breaks (DSBs) in ribosomal DNA (rDNA) repeats. TOPBP1 is recruited to the nucleoli in response to rDNA breaks, and this recruitment is necessary for the inhibition of ribosomal RNA synthesis and nucleolar segregation. This suggests that TOPBP1 plays a key role in the DNA damage response pathway.

Given the role of the DNA damage response pathway in the development of melanoma and the involvement of TOPBP1 in this pathway, it is plausible that the two gene sets have a high overlap because they are both related to the DNA damage response pathway. This could explain the connection between the disease melanoma and the gene TOPBP1, despite the dissimilar abstracts of the two papers. The overlap could be due to the presence of genes involved in the DNA damage response pathway, which is implicated in both the development of melanoma and the function of TOPBP1.
